# Receptor agonist of NFκB signaling ligand directs lung epithelial cell expansion through RANK signaling

**DOI:** 10.1101/2025.05.13.648716

**Authors:** Habibie Habibie, Jelmer R. Vlasma, Kurnia S.S. Putri, Shanshan Song, Marina H. de Jager, Arjen Petersen, Carian E Boorsma, Robbert H Cool, Wim Quax, Martijn C Nawijn, Wim Timens, Janette K. Burgess, Corry-Anke Brandsma, Barbro N Melgert

**Author notes:** Correspondence should be addressed to: Prof. Dr. Barbro N. Melgert. Department of Molecular Pharmacology, Antonius Deusinglaan 1, 9713 AV Groningen, The Netherlands. Contributed equally (Habibie Habibie, Jelmer R. Vlasma).

## Abstract

Chronic obstructive pulmonary disease (COPD) is the third leading cause of death globally, with progressive emphysema driven by repeated epithelial damage and impaired repair. Recently, we found that secretion of the osteokine “receptor agonist of nuclear factor κB signaling ligand” (RANKL) is higher in lung fibroblasts from patients with COPD compared to control and that RANKL represses lung epithelial cell death. However, the underlying mechanism, its cross-species conservation and the involved epithelial cell types remain unclear.

To investigate how RANKL affects lung epithelial cells, we used primary lung organoids from human and mouse epithelial cells and showed that RANKL supplementation increased alveolar organoid forming capacity in these cultures compared to vehicle-treated controls. In elastase-treated mice, RANKL was able to rescue the elastase-induced loss of epithelial cells compared to vehicle-treated controls and it augmented the proportion of epithelial cells in transitional states expressing *Krt8* and *MHCII*. Using A549 epithelial cells to investigate whether RANK or the alternative receptor for RANKL, “leucine-rich repeat-containing G-protein coupled receptor 4” (LGR4) were responding to RANKL treatment, we found that RANKL most likely acts through RANK

These results suggest that RANKL may enhance stem cell survival of primarily alveolar epithelial cells in both mice and humans. We therefore conclude that RANKL is another osteokine, in addition to periostin, osteopontin, osteoglycin, and osteoprotegerin, that has a role in lung tissue repair and that its signaling pathway could be explored for therapeutic applications.

**New & Noteworthy:** RANKL enhances human and murine alveolar epithelial cell expansion. Using organoids and an elastase injury model, we show RANKL promotes alveolar epithelial type II cell expansion and rescues transitional cell type loss. RANKL/RANK signaling therefore emerges as a conserved lung regenerative pathway and may be a potential therapeutic target for diseases like COPD with impaired epithelial repair.

## INTRODUCTION

Chronic obstructive pulmonary disease (COPD) is a prevalent lung condition that currently is the third leading cause of death worldwide^1^. Symptoms of COPD include chronic cough and shortness of breath^1^. In lung tissue, COPD is characterized by widespread alveolar cell damage, manifesting as emphysema, as well as chronic inflammation in the airways^1^. Dysfunctional repair impairs regeneration of lost tissue, presenting as enlargement of alveoli and a loss of lung function^2–4^. In addition to lung damage, patients with COPD are often diagnosed with comorbidities such as osteoporosis and muscle degeneration^5,6^. The receptor activator of nuclear factor-κB (RANK) signaling pathway is believed to drive this accelerated bone degradation and muscle degeneration through interactions with local cell types^7^, activating RANK signaling and promoting tissue degradation. RANKL, the osteokine ligand for RANK, has also been shown to have proliferative effects in lymphoid and breast tissues, particularly on epithelial cells^8–11^. Our recent findings indicate that lung fibroblasts derived from patients with COPD secrete elevated levels of RANKL compared to those from individuals with normal lung function, suggesting a potential role for RANKL in lung epithelial repair processes^12^. This positions RANKL as a candidate osteokine warranting further investigation in the context of lung tissue repair, joining other osteokines such as periostin^13^, osteopontin^14^, osteoglycin^15^, and osteoprotegerin^16^ that have roles in lung tissue.

Understanding the mechanisms of lung epithelial damage and repair is pivotal for identifying factors that could help alleviate the widespread tissue damage observed in COPD. In patients with COPD, emphysema results from loss of alveolar septa and type 1 alveolar epithelial (AT1) cells due to damage and dysfunctional repair responses^17,18^. Upon alveolar damage, AT2 cells are the first cells to respond with proliferation and differentiation into AT1 cells through a pre-AT1 transitional state characterized by Krt8 expression^3,19,20^. In COPD, however, AT2 cells have reduced differentiation capacity as well as elevated necroptosis^21^, indicating defects within this compartment. Repair responses of alveolar epithelial cells have been meticulously characterized in *in vitro models* such as precision-cut lung slices and lung epithelial organoid cultures^22,23^. While insight in the effects of RANKL on alveolar repair is lacking, we previously showed that RANKL can reduce epithelial cell death in precision-cut lung slices derived from mice^24^, indicating a protective role of RANKL in epithelial cells.

In line with its proliferative effects in other epithelial cell types and protective effects in precision-cut lung slices, we hypothesized that RANKL may induce AT2 cell expansion upon alveolar damage enhancing lung alveolar repair. To test this hypothesis, we examined the effect of RANKL in murine and human organoids and performed *in vivo* experiments in a murine model of elastase-induced alveolar lung damage. In addition, to investigate whether RANKL affects AT2 cell differentiation we measured the proportions of various epithelial cell types in control and elastase treated mice treated with RANKL or not. Results of this study identified RANKL signaling as a novel pathway involved in lung epithelial cell expansion and/or differentiation that may be targetable in diseases such as COPD.

## Materials and methods

### Production and purification of soluble RANKL protein

Recombinant soluble RANKL (sRANKL) for in vitro experiments was produced using standard procedures at our facility^25^(*Supplemental Data*). Protein purity was checked by SDS-PAGE and RANKL concentrations were measured with a Coomassie Bradford protein assay kit (#23200, Thermo Fisher) using bovine serum albumin as reference and with a RANKL ELISA kit according to the manufacturer’s instructions (#DY626, R&D Systems, Minneapolis, Minnesota, USA). Values from both assays were always in the same range and the concentration obtained from the ELISA analysis was used for further experiments. Purified sRANKL was routinely tested for endotoxin contamination comparing heat inactivated sRANKL with native sRANKL using RAW 264.7 macrophages and no measurable contamination was found.

### Animal experiments

C57BL6 mice were obtained from Charles River and were maintained with permanent access to food and water in a temperature-controlled environment with a 12h dark/light cycle regimen. Mice were allowed to acclimatize for 7 days prior to experiments. Mouse experiments were performed in the animal facility of the University of Groningen according to strict governmental and international guidelines on animal experimentation. All experiments were approved by the national Care and Use Committee (Centrale Commissie Dierproeven). Female C57BL/6 mice were used in experiments with elastase-induced lung injury (license number AVD10500202011285). Female and male C57BL/6N mice were used for the preparation of murine organoids (license number AVD10500202011285).

### Emphysema-induced lung injury in mice

Female C57BL/6 mice were anaesthetized using 4 % isoflurane, after which 40 U/kg bodyweight porcine pancreatic elastase (Merck, #324682) was intratracheally administered in 40 µl phosphate-buffered saline (PBS). Seven days after the administration of elastase, RANKL treatment was started 3 times per week for two weeks. For RANKL treatment, mice were intranasally exposed to 5 µg RANKL (obtained from Thermo Fisher, #315-11). After two weeks, mice were euthanized by exsanguination through cardiac puncture. The right lung was used for isolation of cells for flow cytometric analysis. The left lung was inflated with PBS-buffered 10% formalin under 20cm H_2_O pressure and fixed overnight at a constant pressure in PBS-buffered 10% formalin and was subsequently embedded in paraffin.

### Flow cytometry

The right lungs of the mice were used for isolation of single cells for flow cytometry. Lungs were minced and incubated in RPMI (#11875093, Gibco, Grand Island, New York, USA) containing 4 mg/ml dispase (#04942086001, Roche, Basel, Switzerland) and 0.1 mg/ml DNAase I (#04716728001, Roche) for 30 minutes at 37 °C. Next, cells were filtered through a 70 µm cell strainer and red blood cells were lysed through osmotic pressure using red blood cell lysis buffer (#00-4333-57, Invitrogen, Carlsbad, California). Cells were resuspended in PFE buffer (PBS + 2 % fetal bovine serum (FBS) + 5 mM EDTA), counted, and one million cells per sample were transferred into a polystyrene tubes. Cells were incubated with Zombie Aqua viability dye (#423101, BioLegend, San Diego, California, USA) in PBS for 20 minutes at room temperature, after which cells were incubated with antibodies targeting extracellular markers EpCAM, MHCII, and Krt8 *(Table S1)*. Samples were analyzed using a Novocyte Quanteon flow cytometer (Agilent, Santa Clara, California). Data were analyzed using *FlowJo* version 10.0 (BD Bioscience, New Jersey, USA). Prior to gating, the plugin PeacQC was used on FSC-A, SSC-A and time parameters to remove aberrant events. We then identified epithelial cells (EpCAM+) within our live cells. Within the epithelial cells, we identified AT2 cells as EpCAM^+^+MHCII^intermediate^, which is similar to previously reported strategies^26^. We also identified Krt8^high^ +EpCAM^high^ cells, previously described as transitional pre-AT1 cells^3,19,20^. Finally, we identified EpCAM^high^ + MHCII^high^ cells as a separate population^27^.

### Cell lines and culture conditions

A549 epithelial cells (ATCC, CCL-185) were cultured in DMEM (#21885025, Gibco), supplemented with 10% FBS, 100 U/ml penicillin/streptomycin (#15140148,Gibco) at 37°C with 5% CO_2_ in humidified air. CCL-206 murine lung fibroblasts were maintained in DMEM (Gibco) / Ham’s F12 medium (#11765054, Gibco) supplemented with 10% FBS, penicillin/streptomycin (100 U/ml) and glutamine (1%, Gibco). MRC5 human lung fibroblasts were maintained in Ham’s F12 medium supplemented with 10% FBS, penicillin/streptomycin (100 U/ml) and glutamine (1%). Prior to organoid culture, CCL206 or MRC5 fibroblast growth was inactivated with mitomycin-C (10 μg/ml, #M4287,Sigma-Aldrich) for 2 h, followed by washing with PBS (Gibco) three times and trypsinization.

### Western blotting

A549 cells were treated with 10 ng/ml RANKL for 15 mins, 30 mins, or 2h. Cell lysates were lysed using ELB-softer (150 nM NaCl, 50 Hepes pH 7.5, 5 mM EDTA, 0.1% NP-40) with one tablet of protease inhibitor cocktail (Thermo Fisher Scientific, Waltham, MA, USA) and one tablet of PhosSTOP phosphatase inhibitor cocktail (Roche, Basal, Switzerland). Protein concentrations were measured using a Pierce BCA protein assay kit (Thermo Fisher Scientific, Waltham, MA, USA) and loaded onto a 10% Bis-Tris gel. Proteins were separated at 100V and subsequently transferred to polyvinylidene fluoride membranes. The membranes were blocked with 5% nonfat dry milk and incubated overnight at 4 °C with one of the following primary antibodies: Akt (1:1000, Cell Signalling), phospho-Akt (1:1000 Cell Signalling) and β-actin (1:10000, Santacruz). The membranes were further incubated with a goat anti-rabbit horseradish-peroxidase-conjugated secondary antibody (1:2000, DAKO) for 1 h at room temperature. For detection, blotted proteins were visualized with an ECLTM Prime Western Blotting System (GE Healthcare). All proteins levels were normalized to β-actin.

### Wnt/β-catenin activity assay

A TOP/FOP flash assay was performed based on a previously published protocol^10^. A549 cells (11000 cells/well) were seeded in 96-well plate. When confluent, cells were transfected with 100 ng/well of either TOP luciferase reporter plasmid or the negative control FOP plasmid using Lipofectamine™ LTX Reagent with PLUS™ Reagent (Invitrogen, #A12621) in serum-free Opti-MEM® medium (Life Technologies, #31985062). After 5 h of transfection, cells were stimulated with either vehicle, positive control CHIR99021 (2 µM), or RANKL (10ng/mL) in Opti-MEM® medium supplemented with 0.1% FBS for 24 hours. After stimulation, cells were lysed using the Bright-Glo™ Luciferase Assay System (Promega, #E2610). Subsequently, luciferase activity was measured using a Synergy HTX Multi-Mode Microplate Reader (BioTek, Winooski, US). Data were collected with Gen5 software (BioTek).

### Human lung tissue

Human lung tissue was used for isolation of epithelial cells for organoid cultures. COPD patients included smoking or ex-smoking individuals with GOLD stages I-IV disease^28^, as defined by degree of airflow obstruction. Characteristics of patients can be found in table S2. Subjects with other lung diseases such as asthma, cystic fibrosis, or interstitial lung diseases were excluded. The study protocol was consistent with the Research Code of the University Medical Center Groningen (research code UMCG (umcgresearch.org) and national ethical and professional guidelines (Code of conduct for Health Research: Gedragscode-Gezondheidsonderzoek-2022.pdf (coreon.org). Lung tissue used in this study was derived from leftover lung material after lung surgery and was exempt from consent in compliance with applicable laws and regulations at the time of experiments (Dutch laws: Medical Treatment Agreement Act (WGBO) art 458 / GDPR art 9/ UAVG art 24). Sections of lung tissue of each patient were stained with a standard hematoxylin and eosin staining and checked for abnormalities by a lung pathologist.

### Cell isolation and organoid culture

Primary lung epithelial cells from mice were isolated and cultured as previously described with modifications^4,23^. The lungs were filled with a dispase (#D4693,BD Biosciences)/agarose (Sigma-Aldrich) mixture and digested at room temperature for 45 min and subsequently homogenized to a single cell suspension. Cells in suspension were negatively selected using a mix of mouse CD45-selecting (#130-052-301, Miltenyi) and mouse CD31-selecting (#130-097-418, Miltenyi) microbeads. Then, CD45-CD31-negative cells were further positively selected with mouse EpCAM-selecting microbeads (Miltenyi, #130-105-958). EpCAM+ cells (10,000) and mitomycin-treated CCL206 fibroblasts (10,000) were resuspended in 100 μl DMEM/F12 medium containing 10% FBS diluted 1:1 with growth-factor-reduced Matrigel (#CLS354230, Corning), and were seeded in a 24-well 0.4 μm transwell insert (Falcon).

Human primary lung epithelial cells were isolated from lung tissue of patients with COPD. Distal human lung tissue was dissociated and homogenized with a multi-tissue dissociation kit 1 (Miltenyi) using a GentleMACS Octo dissociator at 37 °C (Miltenyi). Cells in the resulting suspension were negatively selected using a mix of human CD45-selecting (Miltenyi, #130-045-801) and human CD31-selecting (#130-091-935, Miltenyi,) microbeads. The CD45-CD31-negative cells were further positively selected with human EpCAM-selecting microbeads (#130-061-101, Miltenyi). EpCAM+ cells (5000) were seeded with mitomycin-treated MRC5 fibroblasts (5,000) in Matrigel.

After solidifying, organoid cultures were maintained in DMEM/F12 medium with 5% (v/v) FBS, 1% insulin-transferrin-selenium (#41400045, GibcoTM), recombinant mouse EGF (0·025 µg/ml, #E5160, Sigma-Aldrich), and bovine pituitary extract (30 µg/ml, Sigma-Aldrich). To prevent dissociation-induced apoptosis, ROCK inhibitor (10µM, Y-27632, TOCRIS) was added for the first 48 h. Organoid cultures from mouse and human lung tissue were treated with RANKL (10ng/ml), RANKL-inhibitor osteoprotegerin (OPG, 40ng/mL, #185-OS-025, R&D), a combination of RANKL and OPG (RANKL 10ng/mL + OPG 40ng/mL incubated 37 °C for 1 hour before addition), or vehicle. All treatments were freshly added to the cultures every two days. Organoid cultures were maintained at 37 °C with 5% CO_2_ in humidifled air. The number of organoids was manually counted and organoid diameter was measured at day 14 of culture using an Olympus IX50 light microscope connected to Cell^A software (Olympus).

### Immunofluorescence analysis of lung organoids

Lung organoids were stained for acetylated α-tubulin and pro-surfactant protein C (SFTPC) to identify ciliated airway epithelial cells and AT2 cells, respectively. Briefly, organoids were fixed with ice-cold acetone/methanol (1:1 v/v) and aspecific antibody binding was blocked with 5% bovine serum albumin (BSA, w/v, Sigma-Aldrich). Organoids were then incubated with primary antibodies against acetylated α-tubulin (Santa Cruz Biotechnology, #6-11B-1) and SFTPC (MERCK, #AB3786) in a 1:200 dilution in PBS with 0·1% (w/v) BSA and 0·1% (v/v) Triton-X100 (Thermo Fisher) at 4 ℃ overnight. Organoids were washed three times with PBS and incubated with the secondary antibodies (1:200, donkey anti-rabbit Alexafluor 488, Invitrogen; 1:200, donkey anti-mouse Alexafluor 568, Invitrogen) for 2 h at room temperature. Organoids were washed with PBS three times and then cut from inserts and transferred onto glass slides with mounting medium containing DAPI (Abcam, #ab104139) and a coverslip. Images were obtained using a Leica DM4000b fluorescence microscope connected to Leica Application Suite software (Leica Microsystems).

### Immunohistochemistry

Paraffin-embedded lung tissue was deparaffinized using xylene, after which antigen retrieval was performed using a 10 mM citrate buffer, pH 6. For a staining with Ki67, cells were permeabilized using Triton X for 10 minutes. Next, we performed a blocking step using 5% BSA and 5% normal mouse serum followed by blocking of endogenous peroxidases. Then, primary antibodies for Ki67 (Abcam #AB15580, 1:500) or pro-SPC (Abcam #AB90716, 1:3000) were added for one hour. Secondary antibodies labeled with HRP were finally added. For staining Ki67, we also added a tertiary HRP-coupled antibody. Staining was visualized using NovaRed (Vector Labs, SK-4800) counterstained with hematoxylin. Stained slides were scanned using a Hamamatsu slide scanner (Hamamatsu Photonics).

Scanned imaged were analyzed using color deconvolution implemented in FIJI^29^. For each mouse, 5 random fields of the parenchymal are of the lungs were analyzed for %area positive staining of the entire region of interest. By dividing the %are positive for staining by the %area positive for hematoxylin staining we corrected for the cell number. Averages of the five random fields per mouse were taken and used for group comparisons. For mean linear intercept (LMI) measurements, morphometry was used to calculate the mean linear intercept per section^30^, which was again averaged per mouse.

### ScRNAseq analysis

scRNAseq data generated by *Choi et al* was retrieved from *GSE144468* and processed using *Seurat* version 4.0.1 with R version 4.0.4^20^. As an initial quality control step, cells with more than 10% mitochondrial RNA were removed. Next, data were normalized and scaled, after which variable features (n=4000) were identified and selected for generation of the nearest neighbor graph. Nearest neighbor analysis was done on the first 20 computed principle components and subsequent clustering was performed with resolution set at 0.3. For the 10 identified clusters, epithelial cells were annotated as described by *Choi et al* in their original analyses. Stromal cells, not analyzed by *Choi et al*, were identified by expression of *COL1A1, VIM* and *FN1,* and further annotated according to the treatment the cells received (e.g. stromal cells control). Within the annotated object, expression of *TNFRSF11A (*coding for RANK) was interrogated per cell type.

### Statistical analyses

Biological replicates were defined as lung tissue from one individual mouse or patient. In case of experiments with cell lines, a biological replicate was considered an experiment done on a separate day with a new passage of cells. Data sets with fewer than eight biological replicates were not considered to be normally distributed and tested with nonparametric tests. Datasets with eight or more biological replicates were evaluated for normality using QQ-plots and when normally distributed tested with parametric tests. Log-transformation was used to attain normality when data were not normally distributed. When still not normally distributed after log-transformation, datasets were tested with nonparametric tests.

When comparing two groups Mann-Whitney U (unpaired data) or Wilcoxon (paired data) were used for nonparametric data, while a Student’s t test was used for parametric data (paired or unpaired). For the comparison of multiple groups, we used a Kruskal Wallis (unpaired data) or Friedman (paired data) test for nonparametric data or one-way ANOVA (paired or unpaired) for parametric data. Western blot data using multiple time points of treatment were tested using a repeated-measures ANOVA irrespective of the normality of the data. The data were analyzed using GraphPad Prism 9 (GraphPad software, San Diego, USA). For the *in vivo* experiments comparing effects of treatment with elastase, RANKL, or their combination, a two-way ANOVA with a Sidak post-hoc test was used to identify the effects of each elastase exposure and RANKL treatment separately and to identify when these two treatments were interacting. p<0.05 was considered significant. P-values of each effect are presented below each panel.

## RESULTS

### RANKL promotes growth of murine and human lung organoids

To assess whether RANKL treatment had effects on primary lung epithelial cells, we treated lung organoids generated from both human and murine epithelial cells with recombinant soluble RANKL. Organoids grown from Epcam^+^ cells isolated from mouse lung formed structures resembling either airways or alveoli (*Figure 1a)*. We confirmed the identity of these two different organoid structures by immunofluorescence staining using acetylated alpha-tubulin (marker of ciliated cells present in airway-like organoids) and pro-surfactant protein C (marker of AT2 cells present in alveoli-like organoids) (*Figure 1b*).

**Figure 1.**
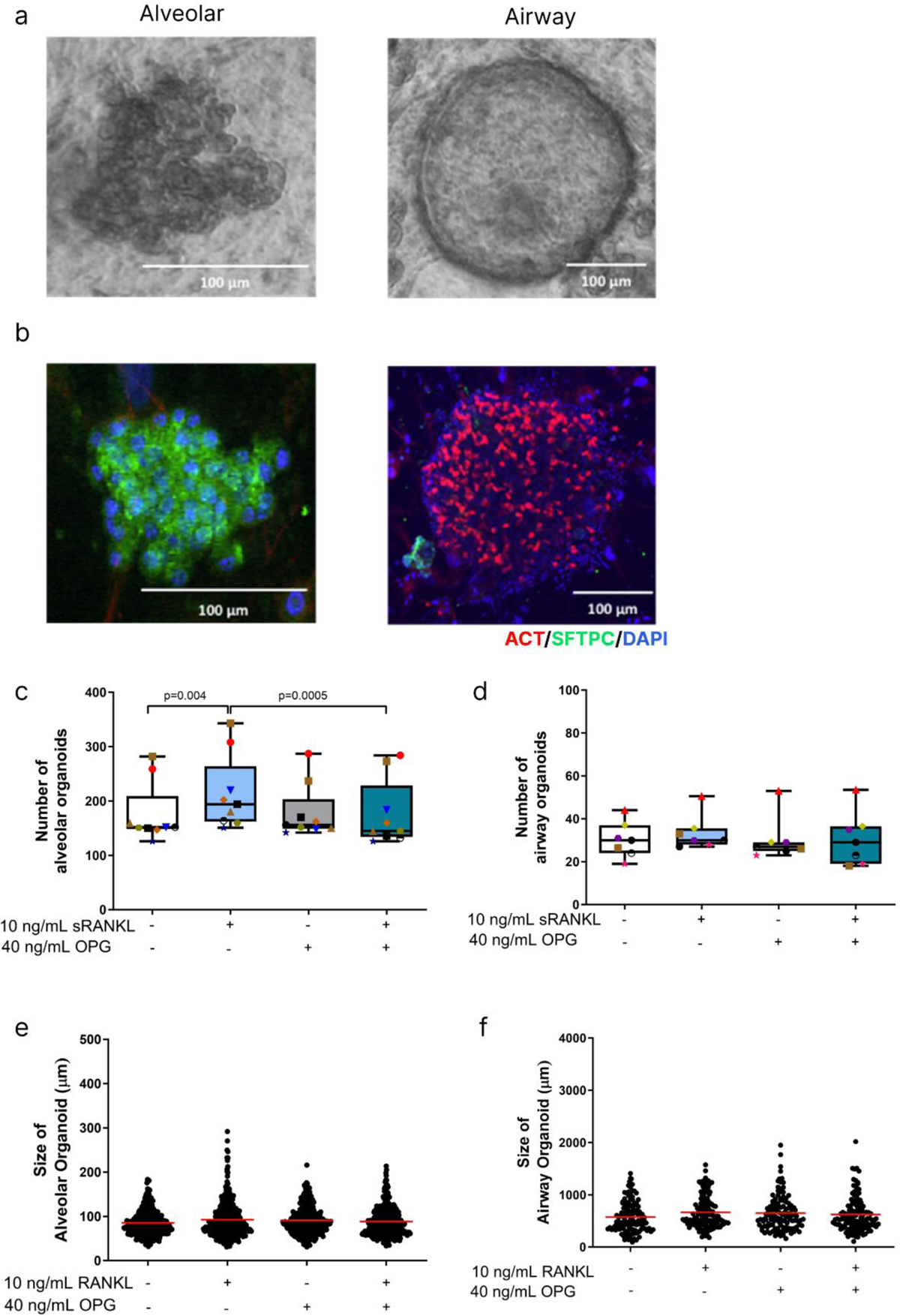
RANKL promotes growth of murine alveolar organoids and this effect is inhibited by osteoprotegerin (OPG). **(a)** Light microscopy images of murine alveolar and airway organoids. **(b)** Immunofluorescence images of murine alveolar and airway organoids stained for acetylated tubulin (ACT, red), pro-surfactant protein C (SFTPC, green), and nuclei (DAPI, blue). Quantification of the number **(c-d)** (each donor marked with different color and symbol) and the size **(e-f)** of murine alveolar organoids and airway organoids after 14 days of incubation with either RANKL (10ng/mL), OPG (40 ng/mL), the combination of RANKL and OPG, or vehicle. Groups were compared using a Friedman multiple comparison test, p<0.05 was considered significant. N=7 biological replicates per group.

RANKL treatment resulted in significantly more alveolar organoids as compared to untreated controls while no effect was seen on airway organoids (*Figure 1c-d*). Notably, co-treatment with OPG, a physiological inhibitor of RANKL, abolished the stimulatory effect of RANKL on alveolar organoid formation. RANKL treatment did not influence the size of either alveolar or airway organoids (*Figure 1e-f*).

To investigate whether these findings in mice could be replicated in primary human epithelial cells, we also generated organoids from Epcam^+^ cells obtained from lung tissue from patients with COPD. These human organoids do not develop into two distinct morphologies as seen for murine organoids but were either mostly alveolar or had a mixed phenotype containing both alveolar and airway epithelial cells (Figure 2a-b). RANKL treatment promoted formation of these alveolar and mixed human lung organoids, which was inhibited in the presence of OPG (Figure 2c). The size of the human lung organoids was not affected by the presence of RANKL (Figure 2d). In summary, treatment of lung organoids derived from primary Epcam^+^ cells with RANKL resulted in a higher number of organoids both in human and murine cultures, an effect that was inhibited by OPG.

**Figure 2.**
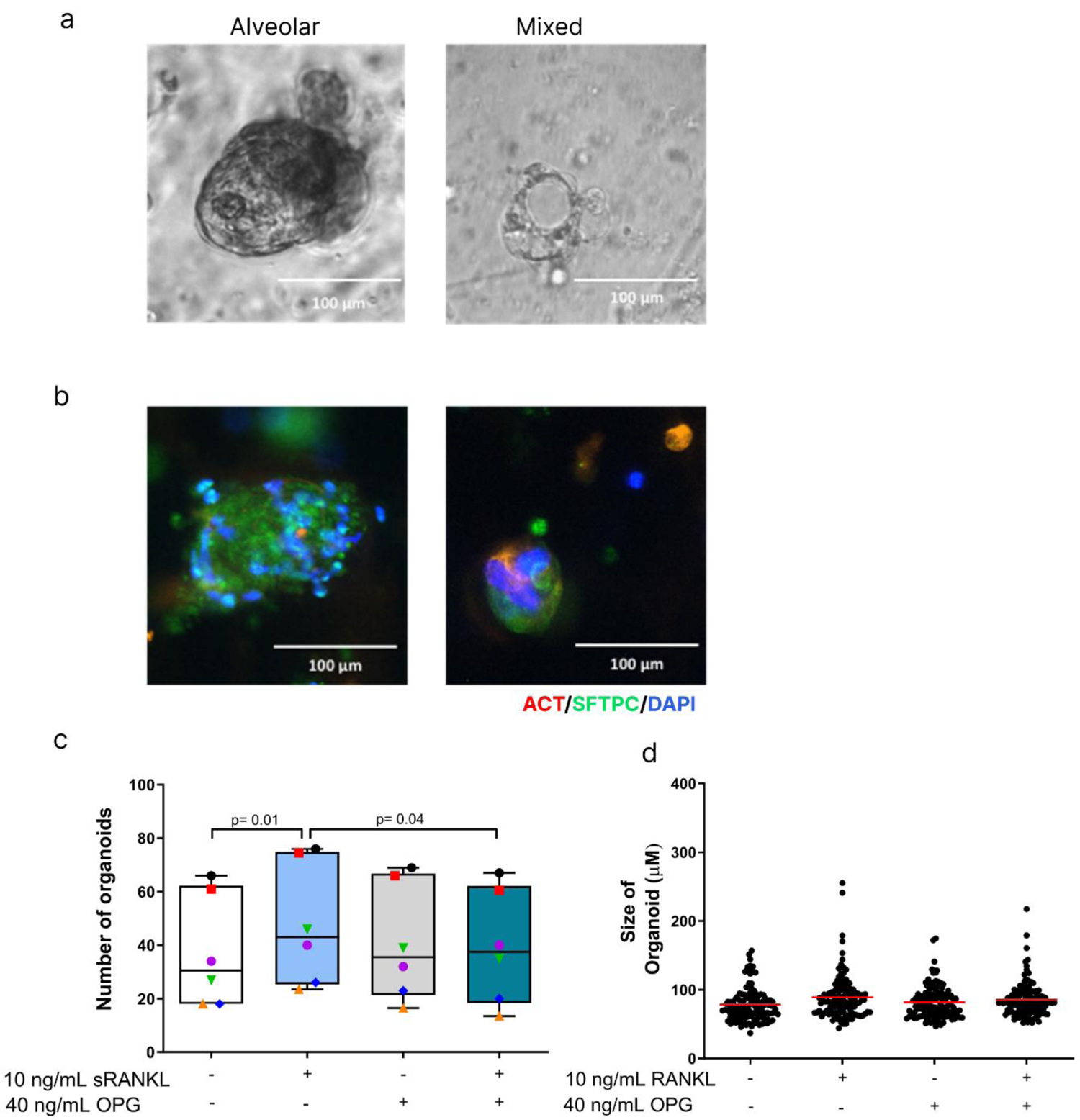
RANKL promotes growth of human lung organoids and this effect is inhibited by osteoprotegerin (OPG). **(a)** Light microscopy image of human lung organoids. **(b)** immunofluorescence image of human lung organoids stained for acetylated tubulin (ACT, red), pro-surfactant protein C (SFTPC, green), and nuclei (DAPI, blue). Quantification of the number **(c)** (each donor marked with different color and symbol) and size **(d)** of human lung organoids after 14 days of incubation with either RANKL (10 ng/mL), OPG (40 ng/mL), the combination of RANKL and OPG, or vehicle. Groups were compared using a Friedman multiple comparison test, p<0.05 was considered significant. N= 6 biological replicates per group.

### RANKL triggers activation of Akt signaling pathway in alveolar type II epithelial cells

To elucidate which receptor signaling was involved in the effect of RANKL on lung organoids, we investigated signaling pathways related to known RANKL receptors, namely RANK and Leucine Rich Repeat Containing G Protein-Coupled Receptor 4 (LGR4)^31^. RANK was previously shown to signal through the Akt pathway^7^, while LGR4 was reported to use the WNT/beta-catenin pathway. We stimulated A549 epithelial cells with RANKL and analyzed both phosphorylation of Akt at several time points by Western blotting and transcriptional activity of beta-catenin with a TOP/FOP dual-luciferase reporter system. Stimulation with RANKL resulted in significant phosphorylation of Akt, 30 minutes after the start of the RANKL stimulation (Figure 3), while no activation of beta-catenin was observed (*Figure S3*). This suggests that RANKL interacts with RANK rather than with LRG4 on A549 cells.

**Figure 3.**
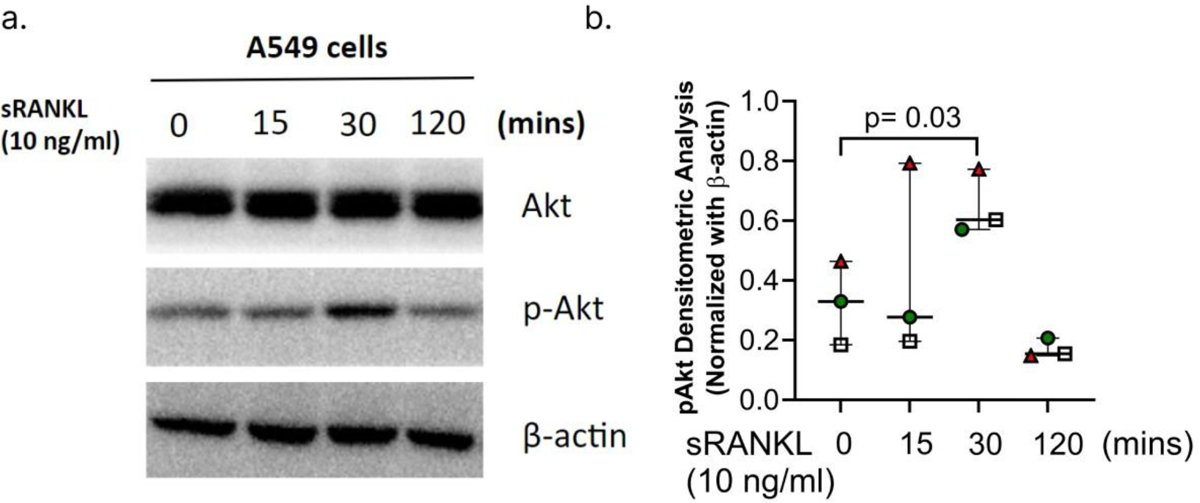
RANKL triggers activation of the Akt signaling pathway in A549 epithelial cells. Representative example of western blot **(a)** and quantification of pAKT activation **(b).** A549 epithelial cells were cultured with 10 ng/mL RANKL and cell lysates were analyzed for phosphorylation of Akt (p-Akt) at several time points by Western blot (n=3). Beta-actin was used as a loading control. Groups were compared using a repeated-measures ANOVA, p<0.05 was considered significant. N=3 biological replicates per group.

### RANKL restores elastase-induced loss of transitional cells

We then investigated which epithelial cell types were involved in the observed RANKL/RANK-mediated organoid expansion. To this end, we characterized the expression of RANK in a previously generated scRNA-seq dataset derived from murine lung organoids by *Choi et a*^20^. Within this dataset, we identified both stromal and epithelial cell types. Expression of RANK *(Tnfrsf11a)* was specific for pre-AT1 transitional cells, a cell state defined by high expression of *KRT8*^3,19,20^.

We next assessed the effects of RANKL on lung epithelial cell proliferation *in vivo* in healthy conditions and after elastase-induced lung injury, as an experimental mouse model for emphysema. To this end, we treated healthy- and elastase-treated mice with saline or RANKL for 2 weeks and assessed proportions of epithelial cells (Figure 5a). After elastase treatment we observed a trend towards a lower proportion of epithelial cells in lung tissue (Figure 5b). RANKL treatment resulted in significantly more *EpCAM^+^* cells as a proportion of total cells (p< 0.02), in line with the results of our organoid experiments.

We then investigated whether the stimulatory effect of RANKL could be specific to certain epithelial cell states observed during the differentiation process of AT2 cells, with a particular focus on pre-AT1 transitional cells that we found to express RANK most highly. To this end, we analyzed AT2 cells (*EpCam^int^, MHCII^int^),* pre-AT1 transitional cells (*Krt8^high^, MHCII^low^, Epcam^high^)* and a subset of AT2 cells characterized by high MHCII expression (*Krt8^high^, MHCII^low^, Epcam^high^) (Figure S1)*. Two-way ANOVA analysis showed that RANKL treatment resulted in significantly more AT2 cells compared to untreated controls *(p<0.001;* Figure 5c). Interestingly, elastase treatment resulted in a significant loss of pre-AT1 transitional cells, while RANKL treatment restored the loss of these cells induced by elastase treatment (Figure 5d). Similarly, RANKL treatment rescued the elastase-induced loss of MHCII^high^ AT2 cells (Figure 5e).

We verified results from our flow cytometry analysis using immunohistochemistry for AT2 cell marker *pro-surfactant protein-C (SFTPC) (*Figure 4f). Though RANKL did not induce more SFTPC staining in control mice, we did observe more SFTPC staining in elastase-treated mice (Figure 4g). we further assessed whether RANKL induced cellular proliferation of alveolar cells by staining for Ki67 (Figure 4h). We found more Ki67 staining in elastase-treated mice compared to control mice, but no effects of RANKL treatment (Figure 4i).

**Figure 4:**
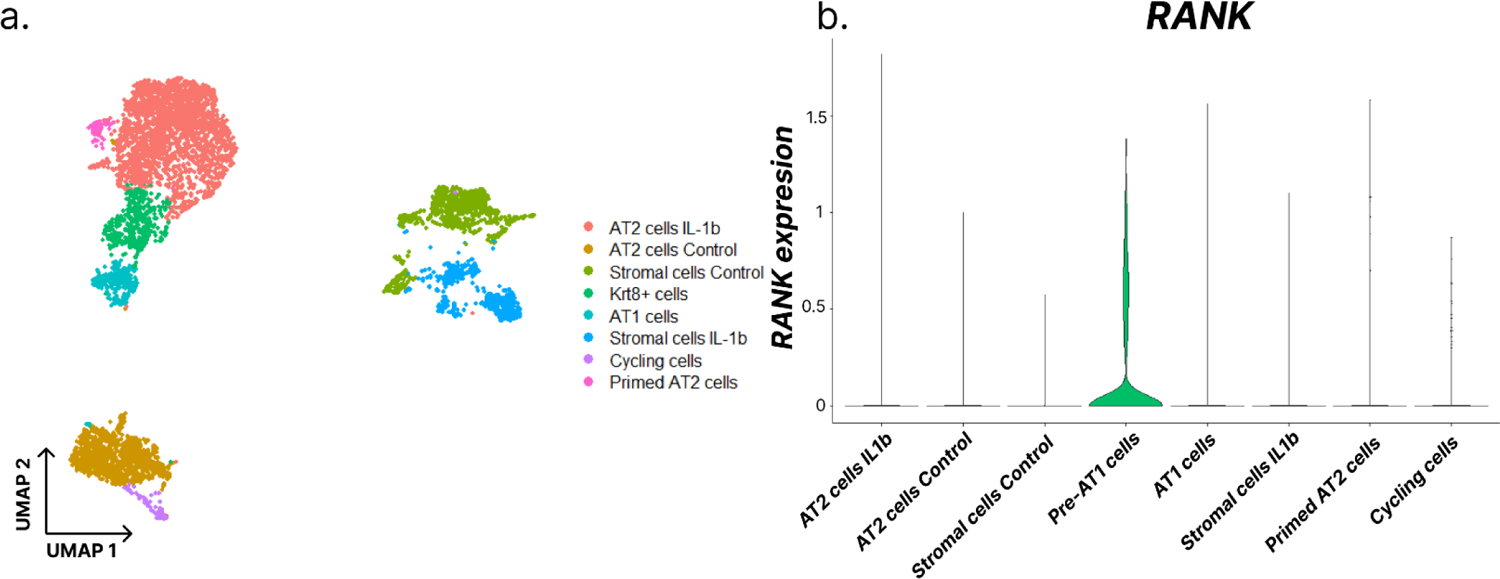
RANK receptor is expressed specifically in pre-AT1 transitional cells. **(a)** UMAP representation of scRNAseq-data generated by Choi et al comprising murine lung organoids. **(b)** Violin plot depicting the expression of RANK in various epithelial and stromal cell types.

**Figure 5.**
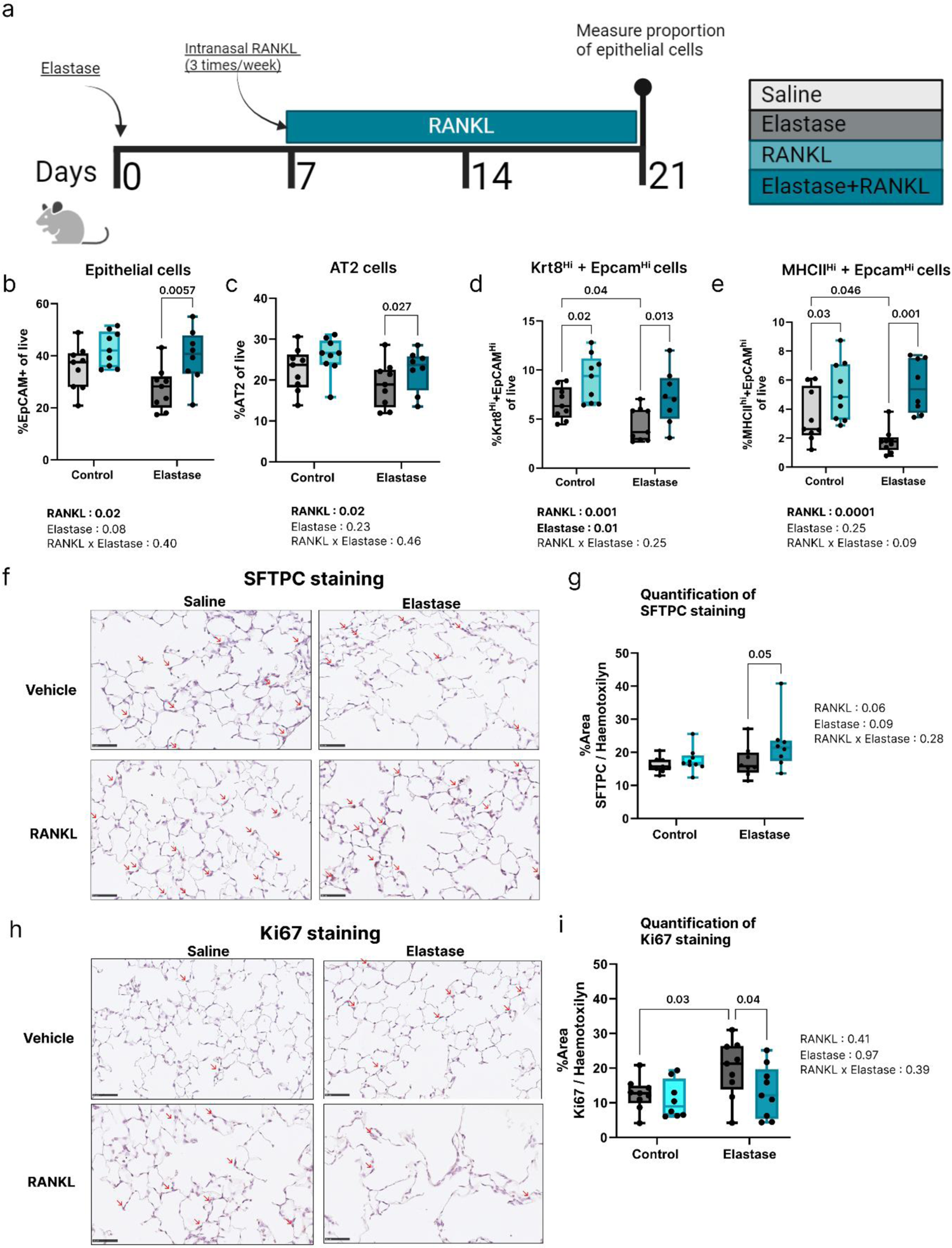
RANKL treatment results in higher proportions of epithelial cells, restoring their loss induced by elastase. **(a)** Experimental scheme describing the setup of this study. Proportions of **(b)** total epithelial cells (EpCAM^+)^, **(c)** type II alveolar cells (AT2), **(d)** MHCII^high^ EpCAM^high^ epithelial cells, and **(e)** Krt8^high^ epithelial cells of all live cells in saline or elastase-treated mice treated with or without RANKL. **(f,g)** Immunohistochemical staining **(f)** and quantification **(g)** of pro-SFTPC in saline or elastase-treated mice treated with or without RANKL. **(h,i)** Immunohistochemical staining **(h)** and quantification **(i)** of pro-Ki67 in saline or elastase-treated mice treated with or without RANKL. Groups were compared using a two-way ANOVA with post-hoc Ŝidák test. Significant P-values of each effect, including the interaction between the effects, are presented below each panel and P<0.05 was considered. N=9 biological replicates per group.

Altogether, these data show a higher proportion of epithelial cells after RANKL treatment, resulting in restoration of elastase-induced loss and elevated proportions of MHCII^high^+EpCAM^High^ and EpCAM^high^+Krt8^Hi^ cells.

## Discussion

COPD is characterized by a progressive loss of alveolar architecture, leading to the development of emphysema. This is thought to be the result of repeated damage combined with dysfunctional repair responses^17,18^. Despite various studies, it is currently unclear which factors can prevent or modify the defects in alveolar epithelial repair observed in COPD. Here, we identified RANK/RANKL signaling as a novel pathway involved in lung epithelial cell expansion, specifically within the alveolar compartment. Using primary lung organoid cultures, we have shown that RANKL treatment resulted in higher alveolar organoid forming-capacity of epithelial cells, implying improved progenitor cell survival and/or proliferation of both human and murine AT2 cells. Furthermore, we demonstrated that RANKL treatment resulted in more Epcam^+^ epithelial cells *in vivo*, both in healthy mice, as well as in those with elastase-induced emphysema, without an effect on cell proliferation. Notably, elastase treatment resulted in loss of MHCII^high^ and KRT8^high^ EpCAM+ cells, indicating a loss of transitional cell types that are important for alveolar repair, which RANKL treatment was able to rescue. Since the mechanism of RANKL-induced epithelial expansion was conserved between mouse and human organoids, this pathway could be of therapeutic interest for diseases characterized by progressive epithelial damage, such as emphysematous COPD.

In organoids and our *in vivo* model, AT2 cells with stem-cell capacity were activated and proliferated to form organoid structures or initiate repair, respectively^22,32^. We observed significant expansion of both MHCII^high^ AT2 and KRT8^high^ epithelial cells *in vivo* after RANKL treatment. Recently, a study by Shenoy *et al.* described MHCII^int^ and MHCII^high^ populations within the AT2 compartment in mice^27^. MHCII^high^ AT2 cells were assigned an important function in interacting with T cells during viral infections^27^. These authors noted that MHCII^high^ cells did not share features with bronchioalveolar stem cells but rather with AT2 cells^19,27^, aligning with our observed outgrowth of alveolar rather than airway structures in murine organoid cultures treated with RANKL. The KRT8^high^ EpCAM state has been modeled to the differentiation trajectory between AT2 and AT1 cells in various murine studies^3,19^. Interestingly, ablation of pre-AT1 transitional state cells after bleomycin-induced lung damage resulted in the development of emphysema, emphasizing their importance during repair^3^. Elastase treatment resulted in a reduction of both MHCII^high^ AT2 and pre-AT1 transitional state cells and interestingly this effect was negated by RANKL treatment. A reduction in the number of AT2 and AT1 cells has been described before for the elastase model as a result of neutrophilic inflammation in response to tissue damage^33^. Considering the anti-apoptotic effects of RANKL we have previously reported^24^, we hypothesize that RANKL rescues alveolar repair through expansion/increased survival of MHCII^high^ AT2 and maybe pre-AT1 transitional state cells. Whether the latter cell type directly responds to RANKL is unclear, but the high expression of RANK we found on these cells suggests they may. Follow-up experiments employing scRNAseq approaches could help to understand how RANKL affects epithelial cell differentiation.

RANKL signaling in epithelial cells resulted in AKT activation, a pathway pivotal for cell growth and survival^34^. Although RANKL was shown to stimulate stromal cell proliferation in secondary lymphoid organs^10^, we recently showed that RANKL also confers protection against cell death in precision-cut lung slices without an effect on cell proliferation, in line with our results on Ki67 activation in this study^24^. Likewise, Benitez et al showed that RANKL-induced signaling results in cell cycle arrest and cell survival through activation of p53 in mammary epithelia^35^. Notably, p53 induction has recently been shown to be a precondition of pre-AT1 transitional cells^3,36^. Therefore, the higher numbers of organoids we found in organoids may be the result of improved survival or activation of AT2 stem-like cells. In COPD, AT2 cells display characteristics of cell cycle arrest and senescence^37–39^, to which the increased RANKL secretion by lung fibroblasts in COPD may be a contributing factor^12^. However, further research is warranted to verify if the anti-apoptotic effects of RANKL also results in a senescent-like phenotype in AT2 cells, or whether it improves cell differentiation into AT1 cells. It therefore remains unclear whether the observed effects of RANKL are beneficial or detrimental to alveolar repair in damaged lungs.

A cross-species conserved RANKL-mediated mechanism of alveolar regeneration could have implications for the treatment of emphysematous COPD. During early stages of COPD, cigarette smoking and viral exacerbations are important causes of alveolar epithelial damage^27^. The influx of neutrophils and the additive oxidative stress results in enhanced epithelial apoptosis as measured by DNA-fragmentation^17^. There is limited data on *in vivo* repair responses in early stages of COPD, but anti-apoptotic effects of RANKL may be beneficial in this process. In this process, stromal cells, previously shown to secrete RANKL in humans^12^, may secrete RANKL to suppress excessive cell death. The elastase model we used is a model of acute lung injury and shows that RANKL treatment is beneficial for epithelial repair. However, in situations of long term constant damage, such as chronic cigarette smoke exposure^40^, the effects of RANKL could be different. Studying whether RANKL supplementation could suppress alveolar damage, and thus stop progression of emphysema, would be a relevant research direction for the treatment of COPD.

There are some limitations to consider in our study. Firstly, in our flow cytometry analysis of lung epithelial cells, we did not include markers for AT1 cells, that can also express MHCII. Nevertheless, characterization of the size and granularity of the MHCII-positive cells indicated that larger AT1 cells were not present among this population^41^, suggesting these are lost during isolation or do not express as much MHCII. Secondly, the induction of AKT signaling by RANKL was demonstrated in A549 cells, which is a ubiquitous cell signaling pathway. Since we only tested two pathways we may have missed alternative pathways through which RANKL can operate. Therefore, these experiments should be validated by exploring other pathways involving AKT. Finally, we investigated how exogenous RANKL affects alveolar proliferation, but did not control for the effects of endogenous RANKL. Future experiments could focus on siRNA applications to investigate whether loss of endogenous RANKL production leads to a reduction in epithelial repair, as this would indicate that endogenous RANKL plays a significant role in an intrinsic repair response in lung tissue.

In conclusion, our results suggest that RANKL may enhance stem cell survival of primarily alveolar epithelial cells in both mice and humans. We therefore conclude that RANKL is another osteokine, in addition to periostin^13^, osteopontin^14^, osteoglycin^15^, and osteoprotegerin^16^, that has a role in lung tissue repair and that its signaling pathway could be explored for therapeutic applications. Based on our data, we suggest that the higher levels of RANKL found in several lung diseases may reflect stimulation of epithelial repair by the lung in an effort to counteract the lung tissue destruction that characterizes these lung diseases.

## Acknowledgements

The funding for this research was partly provided by the Indonesian Endowment Fund for Education (LPDP Scholarship), which supported H.H in pursuing his PhD studies at the Groningen Research Institute of Pharmacy, University of Groningen in the Netherlands. The Noordelijke CARA stichting is also gratefully acknowledged for funding part of the work described. J.R.V. acknowledges support from the Graduate School of Medical Sciences, University of Groningen.

## Acknowledgements

The authors have declared no competing interest.

## Author contributions

HH, JRV, MCN, WT, JKB, CAB and BNM were involved in the conceptualization; HH, JRV, AP, RHC, WQ designed methodology; HH, JRV, KSSP, SS, MHJ, AP, CEB performed experiments and generated data; HH, JRV, and BNM analyzed data; HH, JRV, BNM wrote the original draft; MCN,WT,JKB,CAB and BNM supervised analyses; all authors read and approved the final manuscript; JRV,HH, BNM acquired funding for the project.

## Supplemental Data

### Production and purification of RANKL protein

Plasmid pET15b-mRANKL, encoding the extracellular domain of murine RANKL (RANKL; amino acids 160-316), was transformed into *Escherichia coli* strain BL21(DE3). A 10 mL overnight culture was used to inoculate 1L 2xYT [16g Bacto Tryptone, 10g Bacto Yeast Extract, 5g NaCl, pH 7.0] medium containing 0.1mg/L ampicillin. Cells were grown at 37°C until the absorbance at 600 nm was 0.5 after which RANKL production was induced by adding 0.1 mM isopropyl β-D-1-thiogalactopyranoside and incubation continued for another 16 hours at 20°C. After harvesting and resuspending wet cells at 1 g/3 ml in buffer A [50 mM MES, pH 5.8, 2 mM dithiothreitol, 10 % glycerol], cells were carefully sonicated on ice. Cell debris was removed by centrifugation for 60 min at 38,000xg and 4°C. Supernatants were loaded on an 8 mL cation exchange column (Source 30S, GE Healthcare, USA). RANKL eluted early from a linear 0-500 mM NaCl gradient in buffer A. Pooled fractions at 4°C were brought to pH 7.5 by adding drops of 1 M NaOH, and to 1.3 M (NH_4_)_2_SO_4_ by slowly adding pulverized salt crystals. The following chromatography steps were performed at 10°C. The protein was loaded on a 5 mL HiTrap phenyl sepharose column (GE Healthcare, USA), pre-equilibrated with buffer B [20 mM sodium phosphate buffer, 2 mM dithiothreitol, 10% glycerol, pH 7.5] containing 1.3 M (NH_4_)_2_SO_4_, and eluted early from a linear 1.3-0 M (NH_4_)_2_SO_4_ gradient in buffer B. Pooled fractions were run on a HiTrap Superdex75 column (GE Healthcare, USA) in 20 mM sodium phosphate buffer NaPi, [NaH_2_PO_4_, Na_2_HPO_4_, 10 % glycerol, pH 7.5].

---

**Figure S1:**
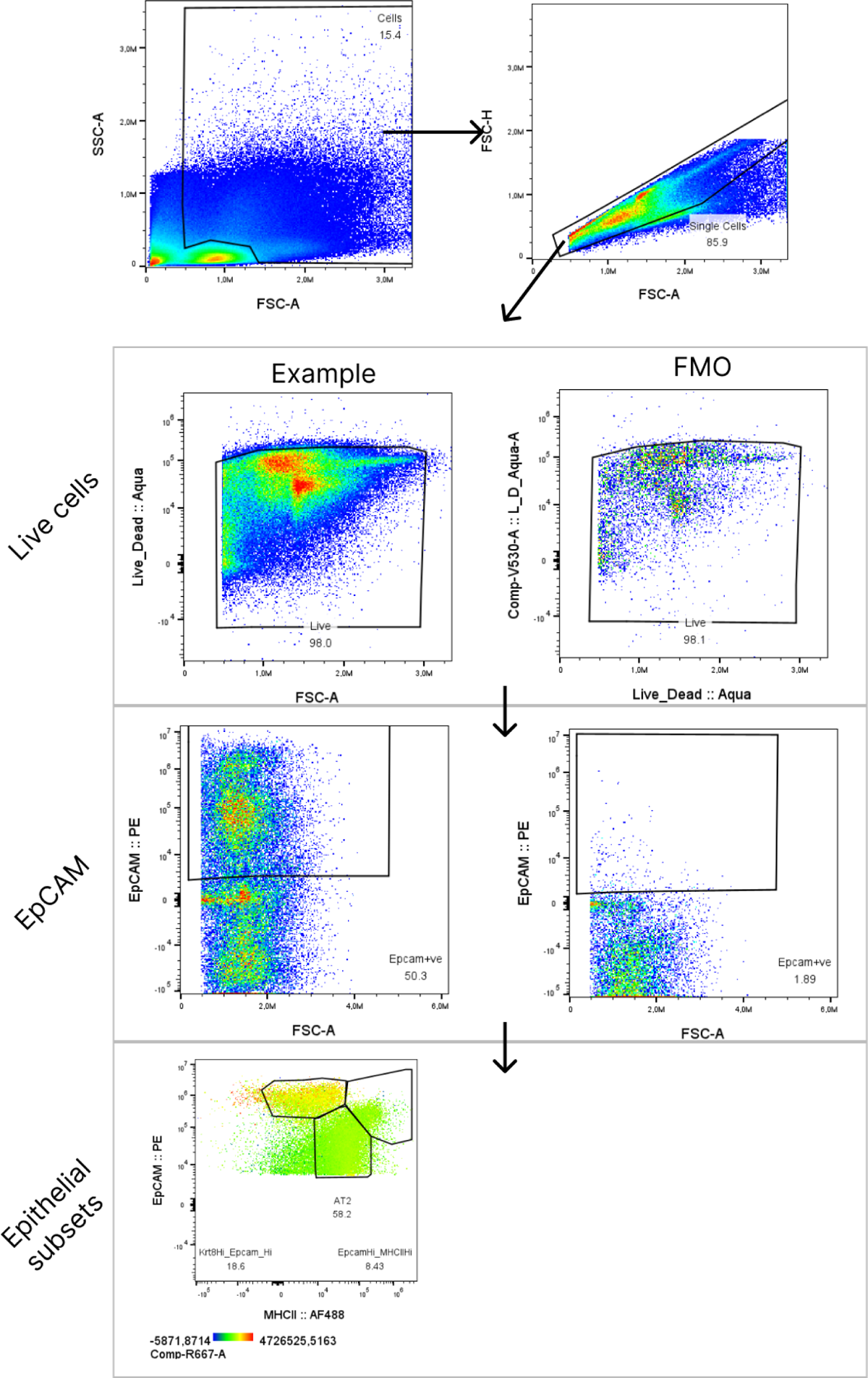
Gating strategy used to identify epithelial subsets in murine lung cells.

**Figure S2.**
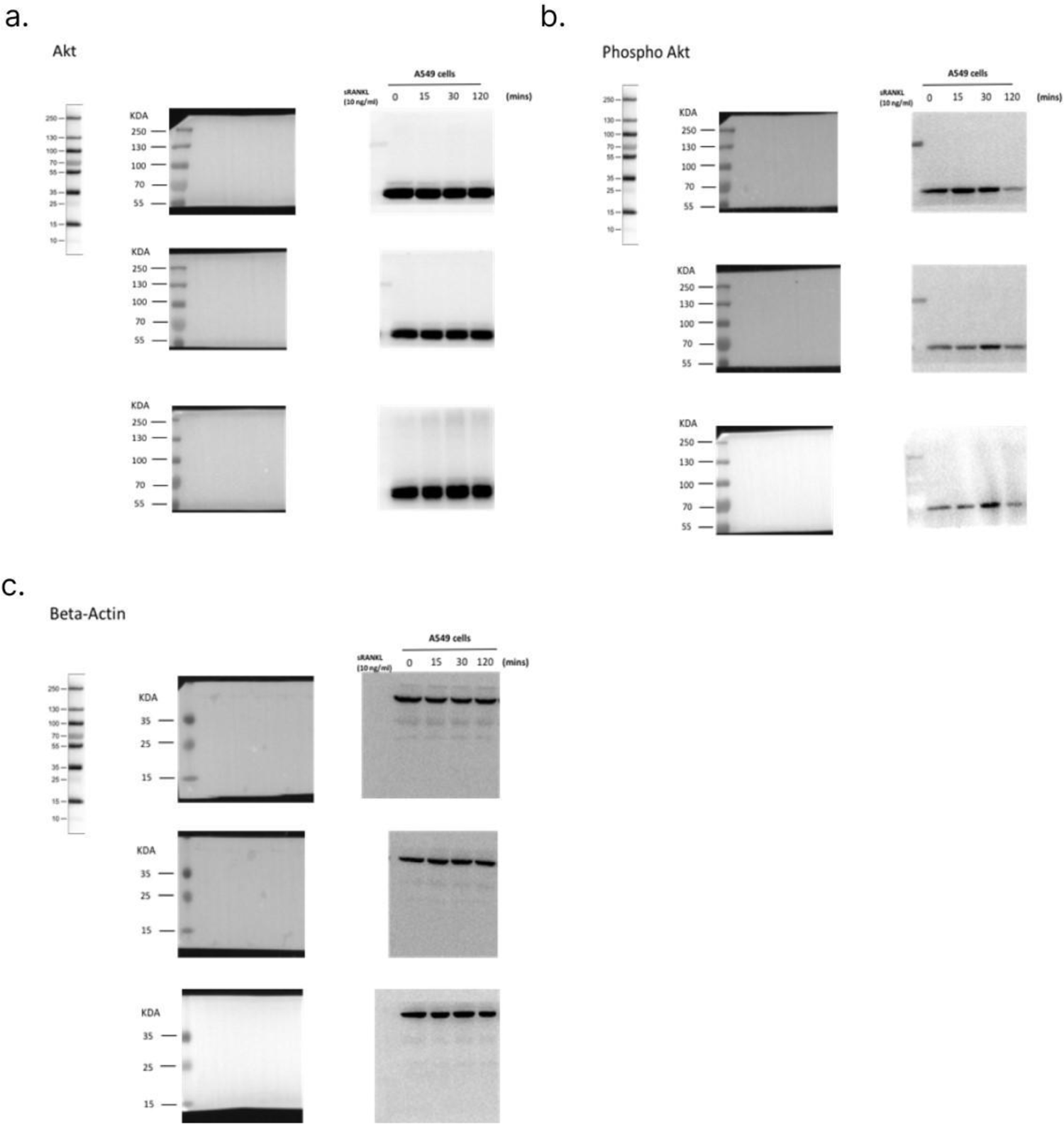
Original western blot data, quantified in figure 3B.

**Figure S3.**
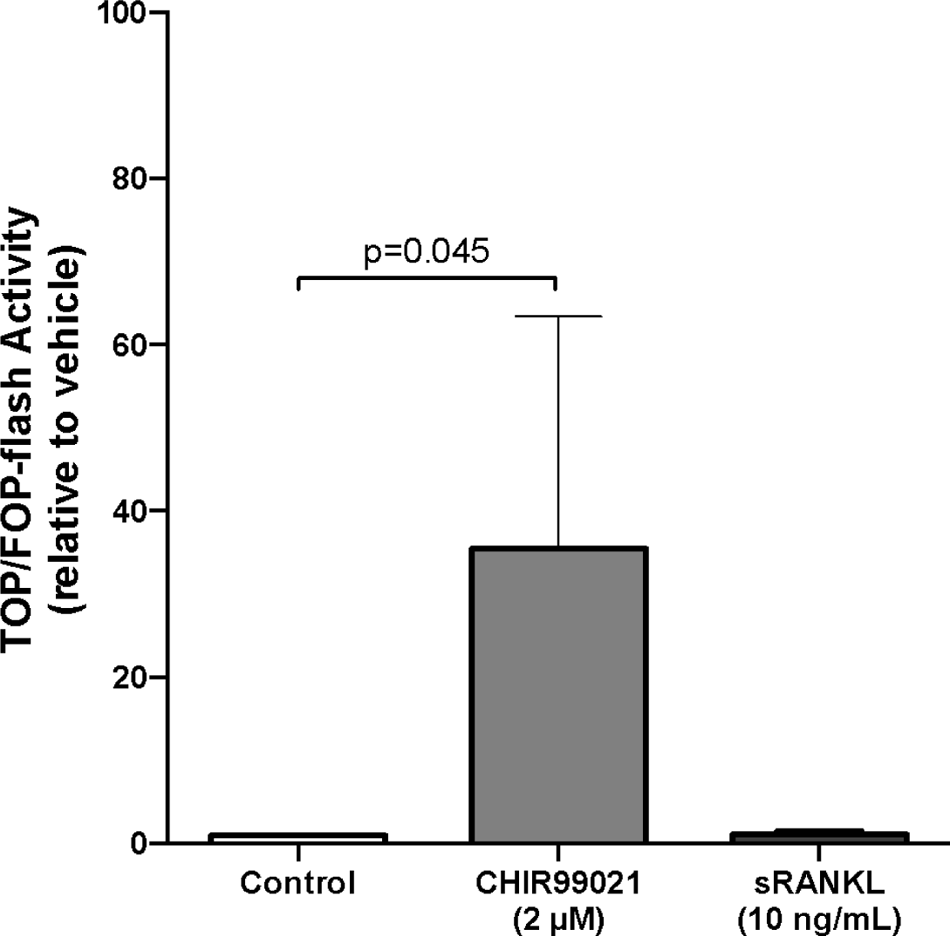
RANKL did not affect the WNT/ beta-catenin signaling pathway in A549 epithelial cells. A549 cells were cultured and transfected with either a TOP or FOP plasmid followed by stimulation with either vehicle, positive control CHIR99021 (2 µM) or RANKL (10ng/mL) for 24 hours (n=3).

**Table S1:**
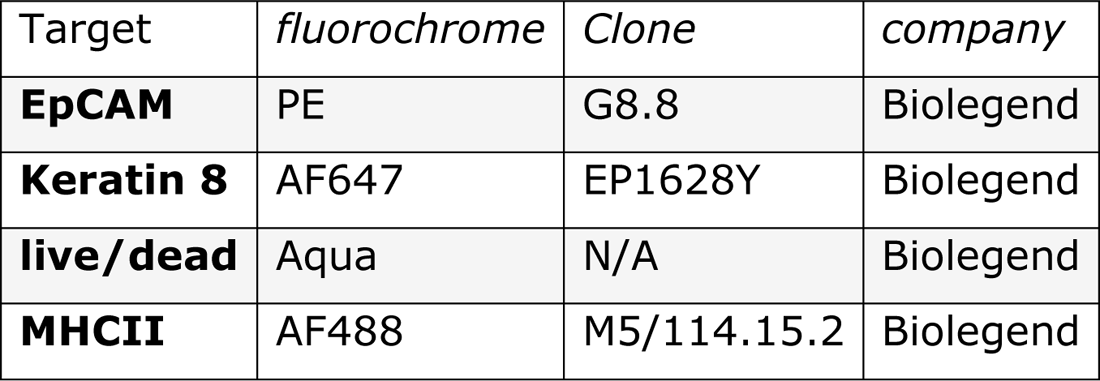
Antibodies used for flowcytometry of murine lung cells.

| Target | *fluorochrome* | *Clone* | *company* |
| --- | --- | --- | --- |
| **EpCAM** | PE | G8.8 | Biolegend |
| **Keratin 8** | AF647 | EP1628Y | Biolegend |
| **live/dead** | Aqua | N/A | Biolegend |
| **MHCII** | AF488 | M5/114.15.2 | Biolegend |

**Table S2.**
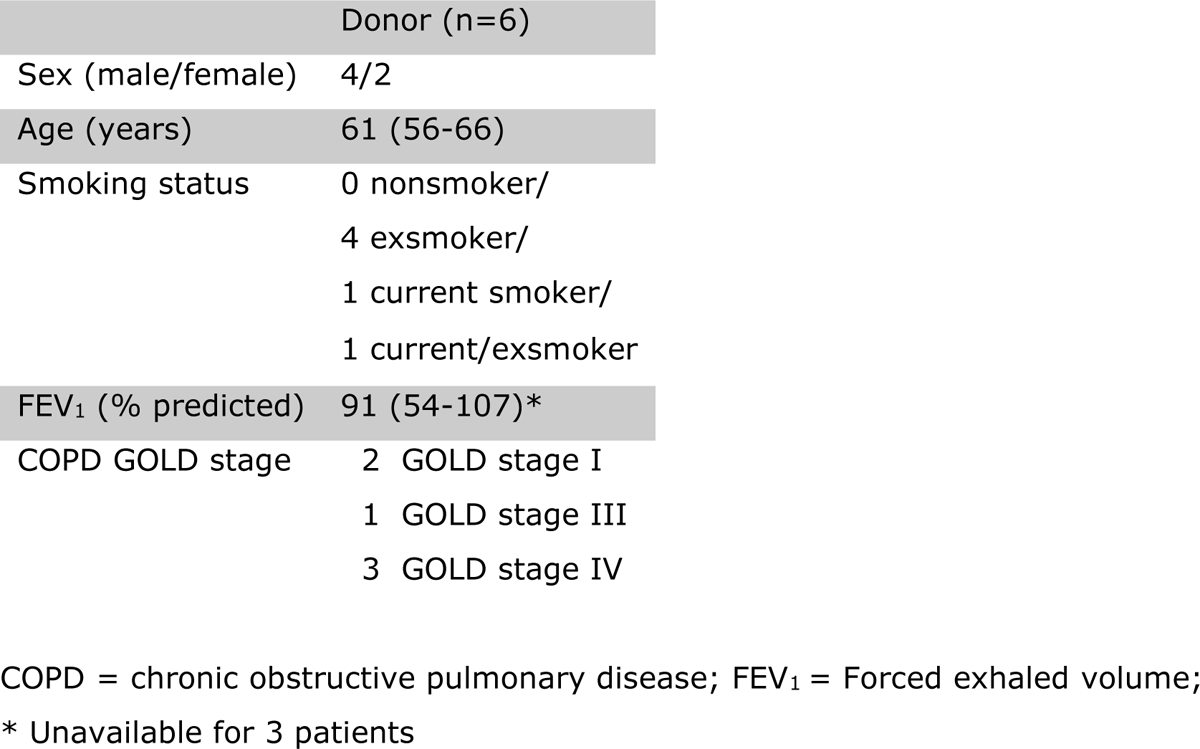
Patient characteristics whose lung epithelial cells were used for organoid cultures. Data presented with median and range.

